# Morc1 re-establishes heterochromatin on activated transposons and shapes the host transcriptome in gonocytes

**DOI:** 10.1101/2023.06.26.546627

**Authors:** Yuta Uneme, Ryu Maeda, Gen Nakayama, Haruka Narita, Naoki Takeda, Ryuji Hiramatsu, Hidenori Nishihara, Ryuichiro Nakato, Yoshiakira Kanai, Kimi Araki, Mikiko C. Siomi, Soichiro Yamanaka

## Abstract

Following reprogramming of DNA methylation, numerous transposable elements (TEs) are transiently activated in male prenatal gonocytes. Persistent expression of such TEs to adulthood leads to arrest of germ cell development. However, how these TEs are re-silenced has been unexplored. Here, we found that DNA-binding protein Morc1 re-established H3K9me3-marked heterochromatin on activated TEs, which involved methyltransferase SetDB1. Although Morc1 also triggered DNA methylation, the types of Morc1-targeted TEs for each epigenetic modification were different, suggesting that these two mechanisms were largely independent of each other. Significant overlap between TEs targeted by Morc1 and those by Miwi2, a nuclear PIWI protein, indicated that piRNA-loaded Miwi2 conferred target specificity to Morc1. Activated TEs drove transcription of adjacent genes by acting as ectopic *cis*-regulatory elements. Such a disrupting effect of TEs on the transcriptome was compensated by Morc1. Thus, Morc1 ensures proper interactions of TEs with host genes through re-establishment of heterochromatin in gonocytes.

## Introduction

In sexually reproducing organisms, germ cells are a specialized group of cells that transmit genetic information to the next generation. Therefore, the genome sequence stored in gametes should be strictly protected from mutagens to maintain the species. Transposable elements (TEs) occupy a substantial portion of the genome in mammals, including humans; percentages of TEs in the genome is much greater than that of protein-coding genes.^1^ Because of their transpositional activity, such genomic parasites constitute a major threat to the integrity of the germline.^2–4^ To counteract such a negative effect of TEs, the host has several silencing systems, one of which is transcriptional silencing by changes in the epigenetic state, such as DNA methylation (DNAme) and histone H3 lysine 9 tri-methylation (H3K9me3).^5–8^

After epigenetic reprogramming of the mouse germline, *de novo* DNAme occurs on whole chromosomes to acquire germ cell identity.^2,9^ This *de novo* DNAme occurs in gonocytes (also known as prospermatogonia) mostly during the embryonic period,^10–13^ playing a critical role in suppressing TE activity.^2,4^ Gonocytes are male germ cells from embryonic day (E) 13.5 to postnatal day (P) 3, representing an essential intermediate between primordial germ cells and spermatogonia in germline development. Dnmt3l, an essential factor for *de novo* DNAme, is expressed during the gonocyte stage. Mice lacking *Dnmt3l* display a hypomethylated genome concomitant with derepression of TEs and severe hypogonadism.^14,15^ Moreover, the proper transcriptome of host genes is disrupted in the mutant,^16^ suggesting the paramount importance to suppress the transpositional activity and ectopic function of TEs as *cis*-regulatory elements.

In animal gonads, TEs are silenced by multiple specialized machineries, one of which is the PIWI-piRNA pathway. PIWI proteins and their associated small non-coding PIWI-interacting RNAs (piRNAs) form the piRNA-induced silencing complexes (piRISCs). They dampen TE activity through post-transcriptional and transcriptional silencing mechanisms.^17–19^ Mice have three PIWIs (Miwi, Mili, and Miwi2), of which Mili and Miwi2 are expressed in gonocyte. MILI degrades cytoplasmic TE transcripts through piRNA-guided endonucleolytic cleavage, producing piRNAs with a complementary sequence to TEs.^20,21^ The resulting piRNAs guide nuclear PIWI protein Miwi2 to active TE loci by tethering the piRISC to the nascent transcript and triggering DNAme.^22–24^ Therefore, piRNAs act as a determinant of target specificity.^25^

Trim28 and KRAB-ZFPs represent another sequence-specific targeting system against TEs in somatic cells and mouse embryonic stem cells (mESCs).^26,27^ KRAB-ZFPs recognize specific DNA sequences of TEs and then recruit SetDB1, a methyltransferase for H3K9, with the assistance of Trim28.^26–30^ Through this series of actions, TEs are enriched with H3K9me3 and eventually silenced at the transcriptional level. Thus, the host triggers enrichment of multiple epigenetic marks, such as DNAme and H3K9me3, depending on the biological context to ensure complete suppression of TEs.

Despite such repression systems, TEs can escape from them and activate in certain developmental stages.^31–37^ We have previously revealed that TEs are transiently upregulated in the middle of the gonocyte stage.^35^ Such TEs tend to cluster on specific genomic regions, termed differentially accessible domains (DADs), which are mostly within intergenic regions.^35^ Together with TE activation, the whole chromosome adopts a relaxed chromatin state to permit *de novo* DNA methyltransferases access to even the heterochromatic region, so that the whole chromosome can undergo DNAme.^35^ However, a sustained active state of TEs beyond the gonocyte stage leads to defects in meiotic prophase, especially double-strand break formation.^38^ These findings strongly suggest the existence of molecular pathways that re-silence TEs in gonocytes. To reveal such a mechanism, we focused on Morc1, a nuclear protein specifically expressed in gonocytes.

Morc1 is a GHKL ATPase that is widely conserved in prokaryotes and eukaryotes.^39,40^ Similar to Miwi2, Morc1 is involved in TE silencing via DNAme,^41^ although how Morc1 triggers repression of TEs remains largely unclear. Here, we revealed that approximately half of the 11,000 transposons that are naturally activated during the gonocyte stage are enriched with H3K9me3, resulting in a transcriptionally inert state as development of gonocytes proceeds. This heterochromatin formation on TEs is largely abolished in *Morc1* knockout (*KO*) mice. Notably, the types of TEs that accumulate DNAme in a Morc1-dependent manner are not the same as those with H3K9me3, suggesting that Morc1 uses different modes of suppression for different types of TEs. Morc1 directly binds to DNA, but it lacks apparent sequence specificity, indicating that other proteins guide Morc1 to its target loci. In this regard, we observed considerable overlap of TEs targeted by Miwi2 and Morc1, suggesting that PIWI-piRNA machinery guides Morc1 to target TEs for heterochromatin formation. In addition to TEs, Morc1 influences the expression of a group of host genes without obvious alteration of the chromatin around their transcriptional start sites (TSSs). Instead, intronic TEs de-repressed in *Morc1 KO* gonocytes coincided with ectopic transcription of specific exons located downstream of such TEs. In summary, Morc1 accumulates the repressive histone modification H3K9me3 on TEs and re-establishes closed chromatin, realizing the proper germline transcriptome built on the relationship between transposons and host genes in gonocytes.

## Results

### Certain TEs activated during the gonocyte stage return to a transcriptionally silent state by accumulation of H3K9me3

In most mammalian tissues, TEs are in a silent state induced by multiple host systems and do not cause genome instability via transpositional activity. However, several reports have revealed that expression of TEs is tolerated in some tissues at specific developmental stages.^31–37^ We previously reported that >10,000 copies of TEs, such as long interspersed nuclear element 1 (LINE-1) and long terminal repeat (LTR) retrotransposons, de-repress in gonocytes of wildtype mice,^35^ while other TEs, including their remnants, remain inactive during this stage (Figure 1A). Such activated transposons in gonocytes turn inactive in the following developmental stage, spermatogonial stem cells,^42^ indicating the existence of specialized machinery that re-silences these elements. To reveal such machinery, we investigated chromatin accessibility at their TSSs during gonocyte development. By comparing read density at ATAC-seq peaks, 14,218 TEs assumed statistically more open chromatin at E16.5 than E13.5. In consistent with a previous report,^35^ more than 60% of such TEs (8,922/14,218) tended to reside in DADs. We termed these activated TEs as DAD TEs. Consistent with the state of chromatin accessibility, DAD TEs produced more transcripts at E16.5 than E13.5 (Figure 1B), then the amount of such transcripts was reduced at P0, suggesting DAD TEs are transcribed in the middle of the gonocyte stage (Figure 1B). Clustering analysis based on the temporal change of chromatin accessibility grouped DAD TEs into four clusters (Figure 1C). DAD TEs in clusters 1 and 2 returned to closed chromatin at P0 in a similar fashion (Figure 1C and 1D), although the extent to which the accessibility reverted to the closed state at P0 was more prominent for TEs in cluster 1. Cluster 3 TEs showed more open chromatin at P0 than earlier stages (Figure 1C and D). Accessibility over cluster 4 TEs at P0 was similar to that at E16.5 (Figure 1C and 1D). Because the overall kinetics of chromatin accessibility was similar in clusters 1 and 2, we categorized them together as Class I TEs. Hereafter, we termed TEs in clusters 3 and 4 as Class II and III TEs, respectively. Class I TEs (6,852; 48.2% of DAD TEs) were re-silenced in gonocytes at P0. Importantly, transcripts from Class I TEs were more significantly reduced compared with those from Class II and III TEs from E16.5 to P0 (Figure 1E), which was in line with their dynamics of chromatin accessibility (Figure 1C). These data indicated that repressive chromatin was re-established on Class I TEs at P0. In public datasets, we found that H3K4me3 and H3K9me3 levels were lower and higher on Class I TEs, respectively, than the other two classes at P0 (Figure 1F). This implied specialized machineries that deposited the repressive histone mark, leading to the formation of closed chromatin on Class I TEs from E16.5 to P0.

**Figure 1.**
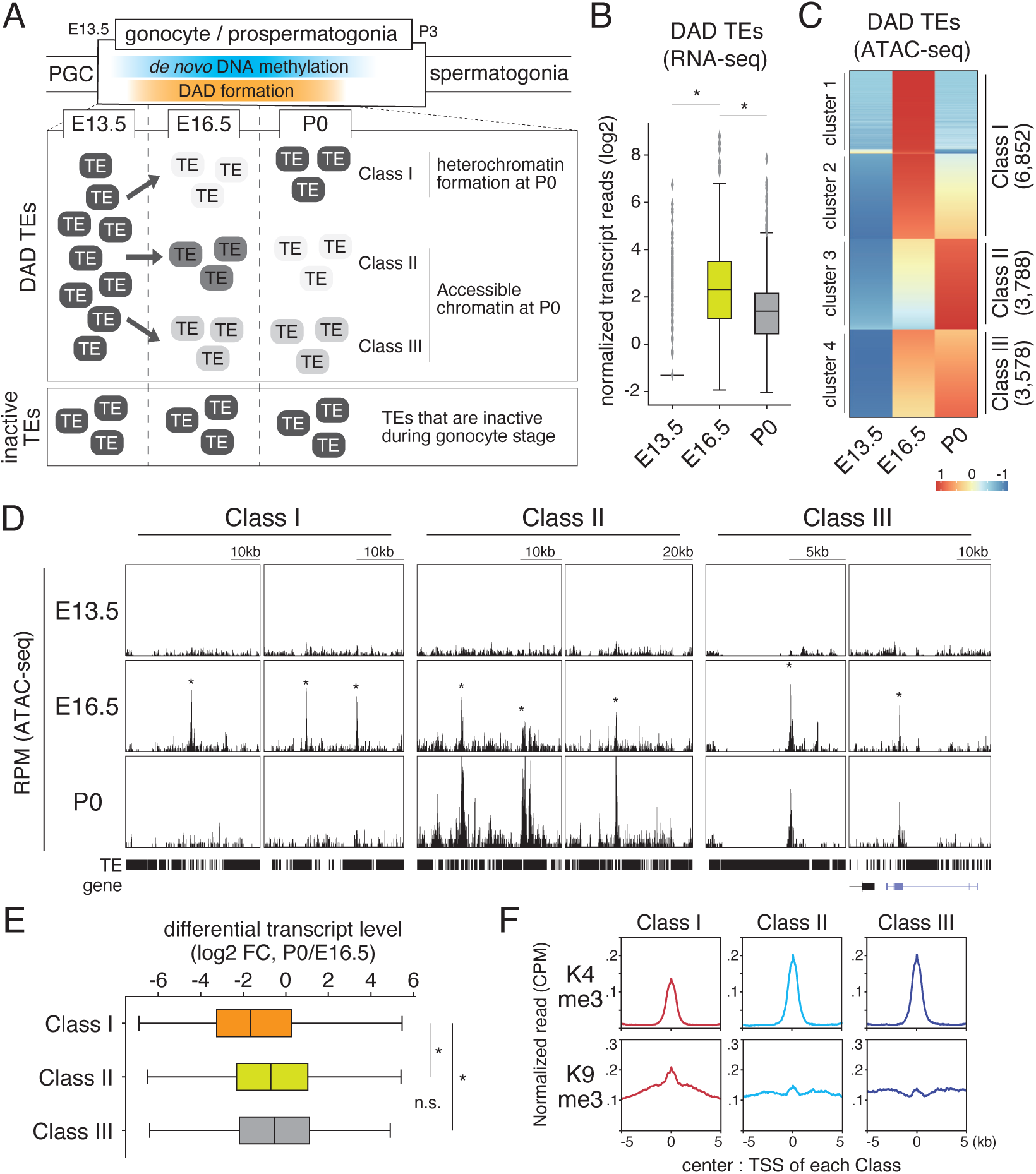
Re-silencing of TEs in gonocytes coincides with enrichment of H3K9me3. (A) Schematic of TE dynamics during the gonocyte stage. TEs are indicated by squares with rounded corners. The color inside the squares denotes the degree of TE silencing. A dark color indicates a more repressive chromatin state on each TE. (B) Dynamics of the transcript abundance of TEs with higher chromatin accessibility at E16.5 than E13.5 during the gonocyte stage. (C) K-mer clustering of DAD TEs based on their dynamics of chromatin accessibility over the TSS region. The number of clusters supplied as input was 4. (D) Genome browser views of representative ATAC-seq peaks categorized to each class. An asterisks marks peaks identified by Homer using ATAC-seq data at E16.5. (E) Differential transcript levels of TEs for each class in gonocytes between E16.5 and P0. The Mann–Whitney U-test was used for statistical hypothesis testing. *p < 0.001. (F) Line plots of the average level of either H3K4me3 or H3K9me3 around the TSS of TEs for each class. Regions surrounding 5 kb upstream and downstream from peaks are depicted.

### Morc1 triggers heterochromatin formation by re-establishing H3K9me3 around Class I TE genomic regions

To reveal such a gonocyte-specific silencing pathway for TEs, we focused on Morc1 for the following reasons. One of the homologue proteins of Morc1 in plants has a physical interaction with histone methyltransferases for H3K9; SUVH2, and SUVH9.^43^ Of note, Morc2a, a MORC family protein in mice, is required for TE silencing in mouse embryonic stem cells (mESCs).^44^ Its human counterpart, Morc2, is also involved in silencing LINE-1.^45^ These findings prompted us to examine whether Morc1 is involved in chromatin compaction and deposition of H3K9me3 on TEs in gonocytes. We performed ATAC-seq at three time points during the gonocyte stage, E13.5, E16.5, and P0, using Morc1 homozygous (*Morc1 KO*) and Morc1 heterozygous (*Morc1 het*) knockout mice. Although there was no clear difference between *Morc1 KO* and *Morc1 het* gonocytes until E16.5, numerous peaks in *Morc1 KO* gonocytes showed higher chromatin accessibility at P0 (Figure 2A). We defined 3,537 peaks that assumed open chromatin in *Morc1 KO* gonocytes at P0 (Figure 2B), 96.9% of which overlapped with TEs such as L1 and LTR retrotransposons (Figure 2C). These results suggested that Morc1 triggered chromatin compaction of TEs during the gonocyte stage. Next, we performed H3K9me3 ChIP-seq at each time point in *Morc1 KO* and *Morc1 het* gonocytes. ChIP-seq reads over 8,491 peaks were significantly reduced in *Morc1 KO* gonocytes at P0 (Figure 2D and 2E). Similar to ATAC-seq, 95.1% of the peaks were on L1 or LTR retrotransposons (Figure 2F). ATAC-seq peaks with higher chromatin accessibility in *Morc1 KO* gonocytes showed a lower signal for H3K9me3 than *Morc1 het* gonocytes (Figure 2G), suggesting its role in chromatin compaction concomitant with deposition of repressive histone marks over TEs.

**Figure 2.**
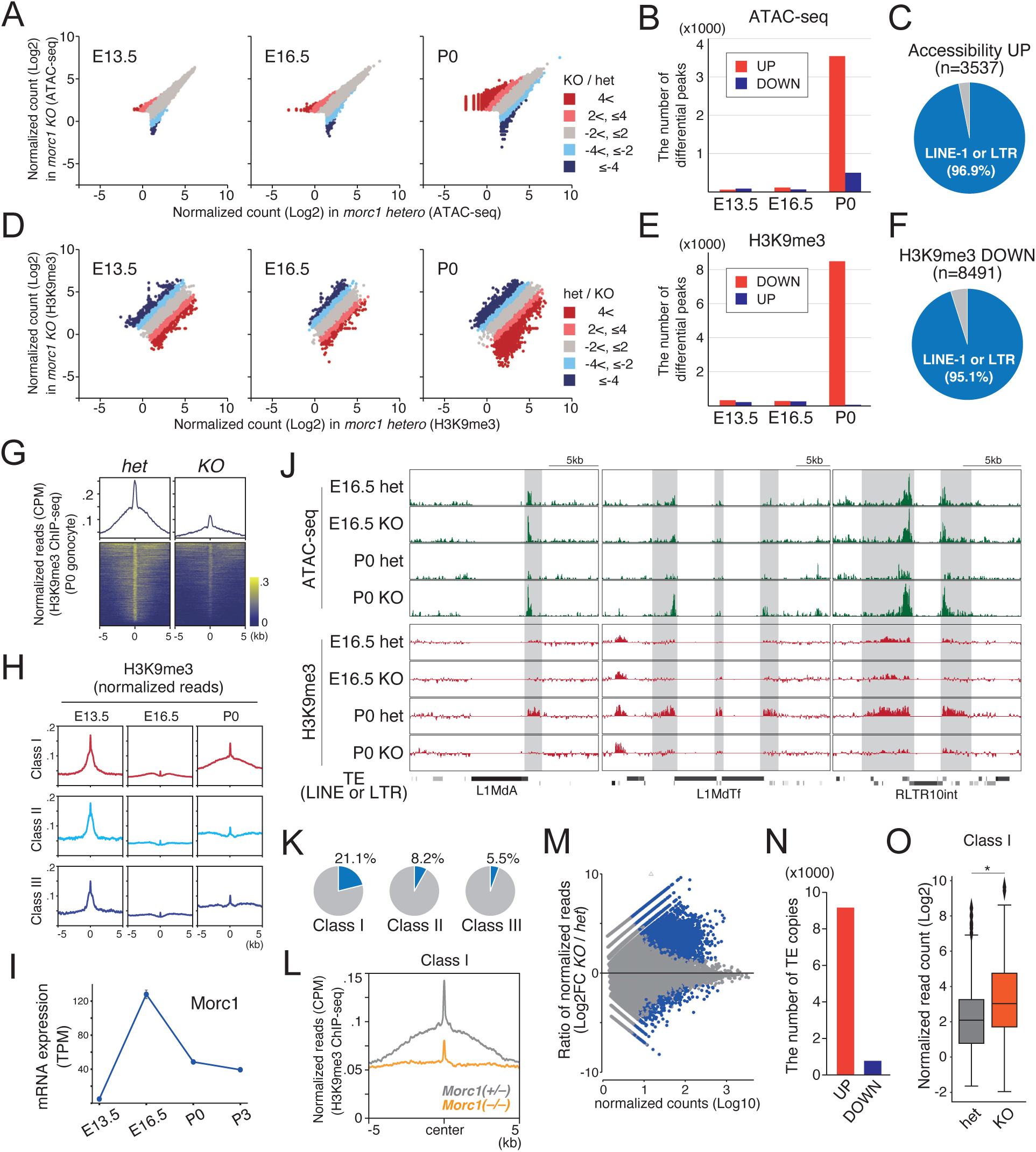
Morc1 triggers the formation of closed chromatin together with enrichment of H3K9me3 over TSSs of Class I TEs. (A) Scatter plots of the normalized number of reads mapped to ATAC-seq peaks for *Morc1 hetero* and *Morc1 KO* gonocytes. The color of each dot indicates the range of the ratio between the two strains. Red, light red, gray, light blue, and blue indicate greater than 4, greater than 2 but no more than 4, greater than −2 but no more than 2, greater than −4 but no more than −2, and no more than −4, respectively. (B) Bar plot of the numbers of peaks that showed differential chromatin accessibility in *Morc1 KO* gonocytes identified by MAnorm. Red and blue bars indicate the number of peaks upregulated and downregulated in *Morc1 KO* gonocytes, respectively. (C) Pie chart of the annotation of ATAC-seq peaks with open chromatin in *Morc1 KO* gonocytes at P0 showing that 96.9% of them overlapped with LINE-1 or LTR-type retrotransposons. (D) Scatter plots of the normalized number of reads mapped to peaks of H3K9me3 for *Morc1 hetero* and *Morc1 KO* gonocytes. The value was normalized to the abundance of H3. The color of each dot indicates the range of the ratio between the two strains. Red, light red, gray, light blue, and blue indicate greater than 4, greater than 2 but no more than 4, greater than −2 but no more than 2, greater than −4 but no more than −2, and no more than −4, respectively. (E) Bar plot of the numbers of peaks that showed a differential level of H3K9me3 in *Morc1 KO* gonocytes identified by MAnorm. Red and blue bars indicate the number of peaks upregulated and downregulated in *Morc1 KO* gonocytes, respectively. (F) Pie chart of the annotation of ChIP-seq peaks with less H3K9me3 signals in *Morc1 KO* gonocytes at P0 showing 95.1% of them overlapped with LINE-1 or LTR-type retrotransposons. (G) Metaplot and heatmap plot analysis of the abundance of H3K9me3 surrounding ATAC-seq peaks that showed significantly higher accessibility in *Morc1 KO* gonocytes compared with *Morc1 hetero* gonocytes at P0. Regions spanning 5 kb upstream and downstream from peak regions are depicted. (H) Temporal transition of H3K9me3 abundance around TSSs of TEs in each class. The H3K9me3 level on class I TEs was more recovered at P0 than that of the other two classes. (I) Expression profile of Morc1 during the gonocyte stage. (J) ATAC-seq and H3K9me3 profiles of *Morc1 hetero* and *Morc1 KO* gonocytes at E16.5 and P0. The positions of LINE-1 and LTR-type retrotransposons from the RepeatMasker database are shown at the bottom. Gray boxes indicate regions showing a difference between *Morc1 hetero* and *Morc1 KO* gonocytes. (K) Pie chart showing the percentages of TEs repressed by Morc1 among TEs in each TE class. (L) As described in (G), but the depicted genomic coordinates were TSSs of Class I TEs. (M) MAplot of RNA-seq reads mapped to TEs in *Morc1 KO* and *Morc1 hetero* gonocytes. Blue dots indicate individual TEs that were differentially expressed between *Morc1 KO* and *Morc1 hetero* gonocytes (FDR < 0.05). (N) Number of TE copies that were significantly upregulated or downregulated in *Morc1 KO* gonocytes compared with *Morc1 hetero* gonocytes (log2 FC >1 or log2 FC <1, FDR < 0.05). (O) Boxplots showing the abundance of transcripts from Class I TEs in *Morc1 KO* and *Morc1 hetero* gonocytes. A t-test was applied for statistical hypothesis testing. *p < 0.001.

All three classes of DAD TEs (Figure 1C and 1D) were enriched with H3K9me3 at E13.5 (Figure 2H) and then lost H3K9me3 from their genomic region at E16.5 upon activation. Because H3K9me3-marked heterochromatin was re-established on Class I TEs during Morc1 expression (Figure 2H and 2I), we suspected that Morc1 was involved in heterochromatin formation on Class I TEs. Visual inspection of some representative genomic insertion sites of Class I TEs revealed that their promoter regions lost H3K9me3 and adopted open chromatin in *Morc1 KO* gonocytes (Figure 2J). Class I TEs included the largest number of Morc1 targets among the three TE classes (Figure 2K). Importantly, re-establishment of H3K9me3 on Class I TEs was severely affected in *Morc1 KO* gonocytes (Figure 2L). RNA-seq analysis revealed that 9,152 TEs were de-repressed in *Morc1 KO* gonocytes at P0 (Figure 2M and 2N). Specifically, Class I TEs were upregulated in the mutant (Figure 2O). These data supported the idea that Morc1 was involved in re-establishment of heterochromatin and silencing of Class I TEs in gonocytes from E16.5 to P0.

### SetDB1 deposits H3K9me3 on Morc1-dependent Class I TEs in gonocytes

Mice express four methyltransferases that target H3K9 residues, SetDB1, Suv39h1, G9a, and GLP.^46–51^ Suv39h1 is responsible for maintenance of H3K9me3 on centromeric repeats.^52,53^ G9a and GLP are involved in converting unmethylated H3K9 to H3K9me2, but not to H3K9me3.^54^ In contrast to these proteins, SetDB1 mediates silencing of TEs through H3K9me3-marked heterochromatin formation in some biological contexts, including mouse embryonic stem cells (mESCs).^28,30^ Therefore, we hypothesized that this enzyme deposited H3K9me3 on Morc1-dependent TEs in gonocytes. To test this hypothesis, we applied an organ culture method to embryonic testes,^55^ because this method enabled evaluation of the effect of a SetDB1 inhibitor (Figure 3A). We chose three genomic sites around TSSs of specific TEs as representative loci targeted by Morc1. Site-specific ChIP-qPCR confirmed that H3K9me3 had accumulated on all three TEs from E16.5 to P0 *in vivo* (Figure 3B). We then examined whether heterochromatin formation on Morc1 targets was recapitulated by the organ culture. Testes extracted from E16.5 embryos were incubated in a culture dish for 3 days, and then germ cells were applied to ChIP analysis (Figure 3A). This revealed that the H3K9me3 level was increased over Morc1-target TEs during the 3 days of *in vitro* culture (Figure 3B). Moreover, adding SETDB1-TTD-IN-1, the SetDB1 inhibitor, led to poor heterochromatin formation on such TEs (Figure 3B). These findings supported the notion that SetDB1 was responsible for catalyzing methylation of H3K9 on Morc1-targeted TEs.

**Figure 3.**
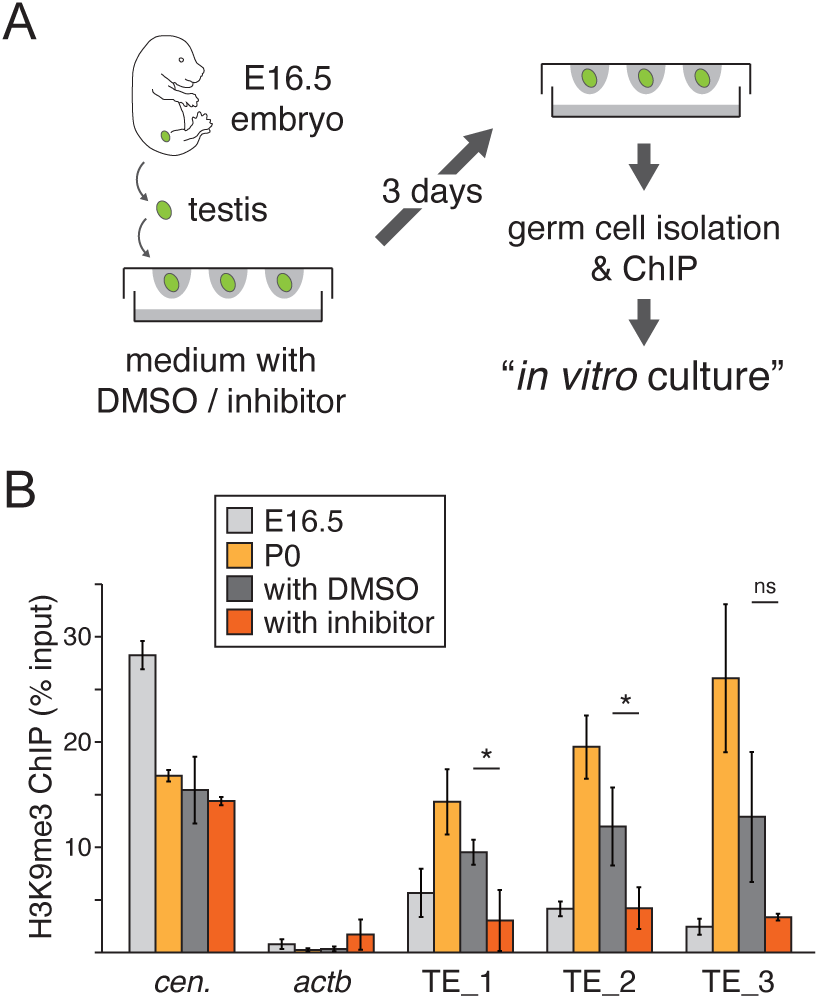
SetDB1 deposits H3K9me3 onto Morc1-dependent TEs in gonocytes. (A) Scheme of gonocyte *in vitro* culture. Testes were excised from E16.5 embryos. The embryo harbored a heterothallic *Ddx4-Venus* transgene, so that germ cells could be isolated by fluorescence. (B) ChIP-qPCR analysis of H3K9me3. The names of four samples used were indicated in the box: E16.5, germ cells isolated from E16.5 testes; P0, germ cells isolated from P0 testis with DMSO: germ cells isolated from testis cultured *in vitro* for 2 days with DMSO, with inhibitor: germ cells isolated from testes cultured *in vitro* for 4 days with a SetDB1 inhibitor. The promoter region of *β-actin* (*actb*) and the pericentromeric region (*cen.*) were used as negative and positive controls, respectively. Three TE regions tested were among Morc1-dependent TEs. A t-test was applied for statistical hypothesis testing. *p < 0.05.

### Target TEs suppressed by H3K9me3 in a Morc1-dependent manner are not equivalent to those suppressed by DNAme

Morc1 triggers accumulation of DNAme on TEs.^41^ Because some reports have shown the hierarchy between H3K9me3 and DNAme in mammalian cell lines,^56,57^ we determined whether such interplay also occurred in Morc1-mediated TE silencing. We compared the families/subfamilies of TEs that gained DNAme [Morc1-dependent TEs (DNAme) or MdTE (DNAme)]^41^ with those that gained H3K9me3 [Morc1-dependent TEs (K9me3) or MdTE (K9me3)]. This revealed that 1,026 TEs out of 8,491 MdTE (K9me3) and 6,302 MdTE (DNAme) were shared between the two groups (Figure 4A). We calculated the proportion of families/subfamilies of TEs of MdTE (K9me3) and MdTE (DNAme) with normalization to the genomic average to estimate which families/subfamilies of TEs were preferentially targeted by Morc1. By sorting MdTE (DNAme) in accordance with the descending order of such enrichment values for MdTE (K9me3), we observed a similar overall pattern of families/subfamilies between MdTE (K9me3) and MdTE (DNAme) (Figure 4B). For example, L1MdTf_I and RLTR10 were the preferential targets for both groups (Figure 4B).

**Figure 4.**
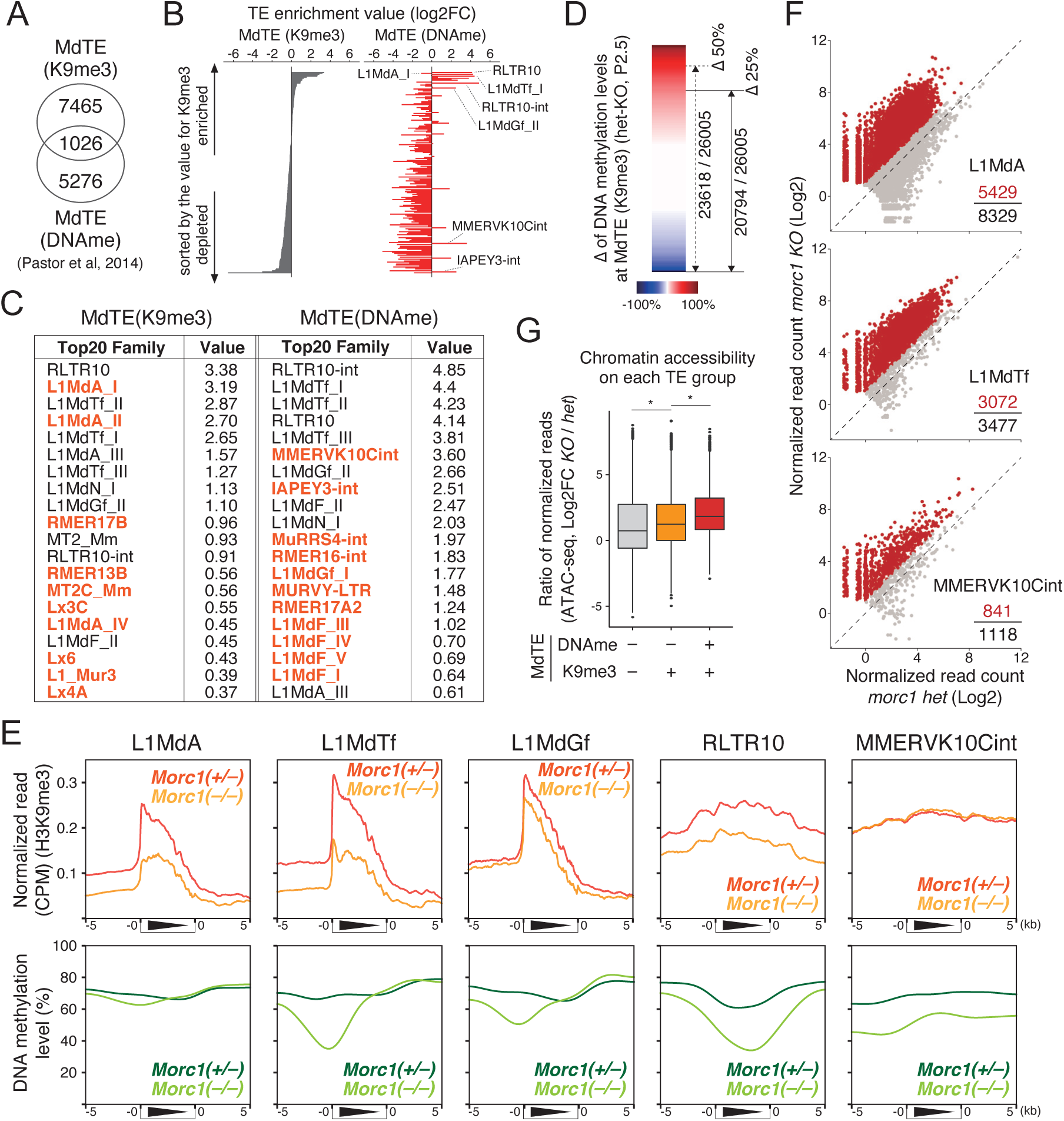
Certain subclasses of TEs acquire H3K9me3 in a Morc1-dependent manner accompanied by little or opposite effect on DNA methylation. (A) Venn diagram showing overlaps between peaks that lost H3K9me3 in *Morc1 KO* gonocytes and differentially methylated regions (DMRs) defined in a previous study.^41^ (B) Superfamilies/subclasses of TEs were sorted by the extent to which certain TEs were enriched in MdTE (K9me3) (see Methods). Enrichment values are also shown for MdTE (DNAme) on the right and sorted by the order of MdTE (K9me3) on the left. (C) Top 20 TE superfamilies/subclasses of MdTE (K9me) and MdTE (DNAme). TEs in red are not shared between the two lists. Values shown were calculated as described in (B). (D) Subtracted difference of the DNAme level between *Morc1 hetero* and *Morc1 KO* gonocytes at MdTE (K9me3). The number of TEs with a value under 50% and 25% are shown. (E) Line plots showing the average level of H3K9me3 (top panels) and DNAme (bottom panels) over regions around each TE superfamily/subclass in *Morc1 KO* and *Morc1 hetero* gonocytes. (F) Scatter plots showing normalized reads of TEs in three TE superfamilies/subclasses in *Morc1 KO* and *Morc1 hetero* gonocytes. (G) Box plots of the ratio of chromatin accessibility between *Morc1 KO* and *Morc1 hetero* gonocytes for three TE categories: TEs that lose H3K9me3 and DNAme in *Morc1 KO* gonocytes, TEs that lose only H3K9me3 in *Morc1 KO* gonocytes, and TEs with no significant change in H3K9me3 or DNAme in *Morc1 KO* gonocytes.

However, there were some distinct differences between them. L1MdA_I ranked second in MdTE (K9me3) was avoided as a target of MdTE (DNAme) (Figure 4B). Conversely, MMERVK10Cint had a negative enrichment value in MdTE (K9me3), although it was the preferential target of Morc1-dependent DNAme. Such a difference was obvious when the top 20 families/subfamilies of each group were compared (Figure 4C). Some subfamilies of L1MdA were included in the top 20 for MdTE (K9me3), but were not in the counterpart of MdTE (DNAme) (Figure 4C), suggesting that Morc1 deposited two different epigenetic marks on different genomic regions. In support of this, 20,794 and 23,618 MdTE (K9me3) out of 26,005 were not included in previously annotated differentially methylated regions (DMRs),^41^ when the threshold of the differential DNAme level between *Morc1 KO* and *Morc1 het* gonocytes was set at 25% and 50%, respectively (Figure 4D). An average line plot of the epigenome marks over each family/subfamily also showed such a trend (Figure 4E). H3K9me3 over promoter regions of L1MdA, L1MdTf, and L1MdGf were reduced in *Morc1 KO* gonocytes (Figure 4E). Among the three L1 subfamilies, the H3K9me3 level on L1MdGf was least affected in *Morc1 KO* gonocytes. Conversely, the degree of DNAme loss over L1MdA was the smallest among them. For LTR retrotransposons, the H3K9me3 level over MMERVK10Cint in *Morc1 KO* gonocytes was almost the same with that in *Morc1 het* gonocytes. Interestingly, the DNAme level over this LTR element was reduced in *Morc1 KO* gonocytes (Figure 4C and 4E), although another LTR element, RLTR10, lost both H3K9me3 and DNAme in *Morc1 KO* gonocytes (Figure 4C and 4E). These results indicated that Morc1 suppressed L1MdA and MMERVK10Cint via H3K9me3 and DNAme, respectively, while L1MdTf was repressed by Morc1 through both epigenomic marks. In terms of transcripts, numerous copies of the three families/subfamilies (L1MdA, L1MdTf, and MMERVK10Cint) were de-repressed in *Morc1 KO* gonocytes, indicating that both H3K9me3 and DNAme suppressed the transcription of these TEs (Figure 4F). Moreover, chromatin accessibility was increased over MdTE (K9me3), which was not categorized as MdTE (DNAme) (Figure 4G), supporting the notion that H3K9me3 alone exerted a repressive effect on TEs. These data suggested certain overlap between MdTE (K9me) and MdTE (DNAme), while some families/subfamilies lost only one of the two epigenetic marks in *Morc1 KO* gonocytes. Thus, Morc1 used different modes of suppression for different types of TEs.

### Miwi2-piRISC is involved in target TE selection for heterochromatin formation by Morc1

More than 90% of Morc1 targets were TEs (Figure 2C and 2F). Similar to Morc1 in worms,^58^ mouse Morc1 binds directly to DNA, although such binding did not show obvious sequence specificity (Supplementary Figure 1A and 1B). These data imply that other factor(s) conferred target specificity to Morc1. We suspected that Miwi2 was involved in such target recognition for the following reasons. Among the three PIWI family proteins in mice, Miwi2 is the only nuclear protein expressed in gonocytes (Figure 5A). Miwi2 is required for enrichment of H3K9me3 in P10 spermatogonia on specific TEs, such as L1MdA and L1MdT,^59^ both of which were among the TEs preferentially targeted by Morc1. These findings prompted us to compare TEs regulated by Morc1 with those regulated by Miwi2 in more detail. By reanalyzing the H3K9me3 ChIP-seq data from *Miwi2 KO* germ cells,^59^ we found substantial overlap of ChIP-seq peaks targeted by either Morc1 or Miwi2 (Figure 5B). The H3K9me3 level over peaks dependent on Miwi2 was decreased in *Morc1 KO* gonocytes and vice versa (Figure 5C). The effect of *Miwi2 KO* around genomic insertion sites of each family/subfamily was similar to that of *Morc1 KO* (Figure 5D), although the distribution patterns of H3K9me3 over TEs were different, possibly because of the difference in ChIP-seq library preparation. piRNAs loaded on Miwi2 were rich in antisense sequences for their target TEs. Therefore, the abundance of mapped piRNAs over a specific TE family/subfamily are useful to measure the degree of dependency on the PIWI-piRNA pathway for its repression. This analysis revealed that the dependency for suppression of Morc1-targeted TEs in the PIWI-piRNA pathway was indistinguishable from that of Miwi2-targeted TEs (Figure 5E). Such a trend was not observed for randomly selected TEs (Figure 5E). Importantly, TEs targeted by Trim28, which mediates TE silencing via KRAB-ZFPs, showed significantly less dependency on the PIWI-piRNA pathway than the others (Figure 5E). Additionally, more than half of DMRs in *Miwi2 KO* gonocytes showed severe loss of DNAme in *Morc1 KO* gonocytes (Figure 5F). Comparing this result with the data in Figure 4D supported the significant similarity of target TEs of Morc1 and Miwi2. These data suggested that Morc1 and Miwi2 shared their target TEs, and supported that Morc1 recognized its targets with the aid of the PIWI-piRNA pathway.

**Figure 5.**
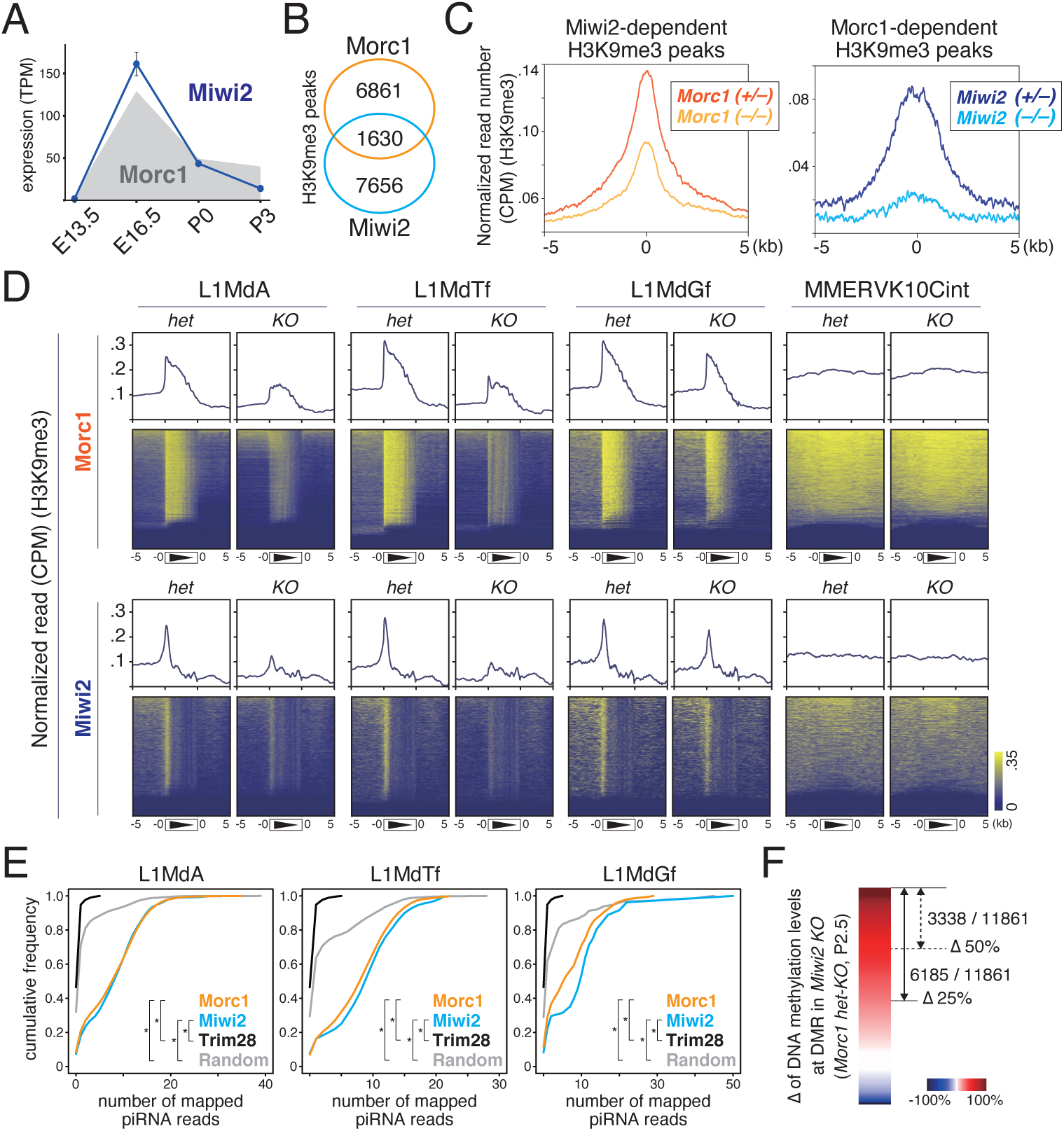
Substantial number of TEs targeted by Morc1 are preferentially repressed by Miwi2. (A) Temporal change of the abundance of Miwi2 transcripts during the gonocyte stage. Shaded gray indicates the pattern of Morc1 expression shown in Figure 2H. (B) Venn diagram showing the overlaps between peaks that lost H3K9me3 in *Morc1 KO* and *Miwi2 KO* gonocytes.^59^ (C) (top panel) Average plots showing the H3K9me3 level in *Morc1 hetero* and *Morc1 KO* gonocytes over H3K9me3 peaks dependent on Miwi2 (left panel). Average plots showing the H3K9me3 level in *Miwi2 hetero* and *Miwi2 KO* gonocytes over H3K9me3 peaks dependent on Morc1 (right panel). (D) Heat map showing the average level of H3K9me3 over regions containing the indicated TE superfamilies/subclasses in *Morc1 KO* and *Morc1 hetero* gonocytes (top panel), and *Miwi2 KO* and *Miwi2 hetero* gonocytes (bottom panel). (E) Cumulative plots showing the distributions of piRNA reads mapped to each copy of the shown TE superfamilies. In addition to the target TEs of Morc1 (orange), Miwi2 (blue), and Trim28 (black), the plots of randomly selected TEs (gray) are also shown. The x-axis shows the number of reads mapped to each TE copy. The y-axis shows the cumulative fraction. The two-sided Kolmogorov–Smirnov test was applied for statistical hypothesis testing. *p < 0.001. (F) As described in Figure 4D, the subtracted difference in the DNAme level between *Morc1 hetero* and *Morc1 KO* gonocytes at DMR in *Miwi2 KO* gonocytes.

### Morc1 compensates the host gene transcriptome that is transiently deregulated during the gonocyte stage

To determine the effect of Morc1 on genomic elements other than TEs, we investigated changes in host gene expression of *Morc1 KO* gonocytes. More upregulated than downregulated genes were found in *Morc1 KO* gonocytes at P0 and P3 (Figure 6A). There were 203 and 323 upregulated genes at P0 and P3, respectively, among which 137 genes were shared between the two stages (Figure 6B and 6C). We assumed that chromatin around TSS regions of the upregulated genes had an open structure in *Morc1 KO* gonocytes, but we could barely see such changes at both stages (Figure 6D). Accordingly, there were little changes in the H3K9me3 level on TSSs (Figure 6E). Genome browser view around the upregulated genes revealed that the reads of RNA-seq were mapped to a part of their exons (faded red boxes in Figure 6F) and not to their TSS regions. Importantly, the neighboring TE (faded blue boxes in Figure 6F) appeared to serve as an alternative promoter. Such TEs lost H3K9me3 and DNAme in *Morc1 KO* gonocytes (Figure 6F). These observations indicated that Morc1-dependent TEs acted as ectopic *cis*-regulatory elements that disrupted the expression of neighboring genes in *Morc1 KO* gonocytes. In support of this, genes upregulated in *Morc1 KO* gonocytes were located significantly closer on the genome *in cis* to the upregulated TEs compared with randomly selected TEs (Figure 6G). Next, using a time course RNA-seq dataset from wildtype gonocytes, we profiled the temporal change in transcript abundance of 137 genes upregulated in *Morc1 KO* gonocytes, revealing that expression of such genes was upregulated at E16.5 and then reduced at P0 in wildtype gonocytes (Figure 6H), which was reminiscent of the dynamics of Class I TEs (Figures 1 and 2). These 137 genes included genes that are not usually expressed in testes, such as Ranbp3l, which is highly expressed in kidneys, and Sult2a5, which is exclusively expressed in the liver.^60^ These results showed that ectopic expression of some genes without a clear function in testes occurred in gonocytes at E16.5 and such expression became repressed in a Morc1-dependent manner at P0. Thus, Morc1 corrected the host gene transcriptome through suppression of nearby TEs that would otherwise disrupt the host gene expression network (Figure 7).

**Figure 6.**
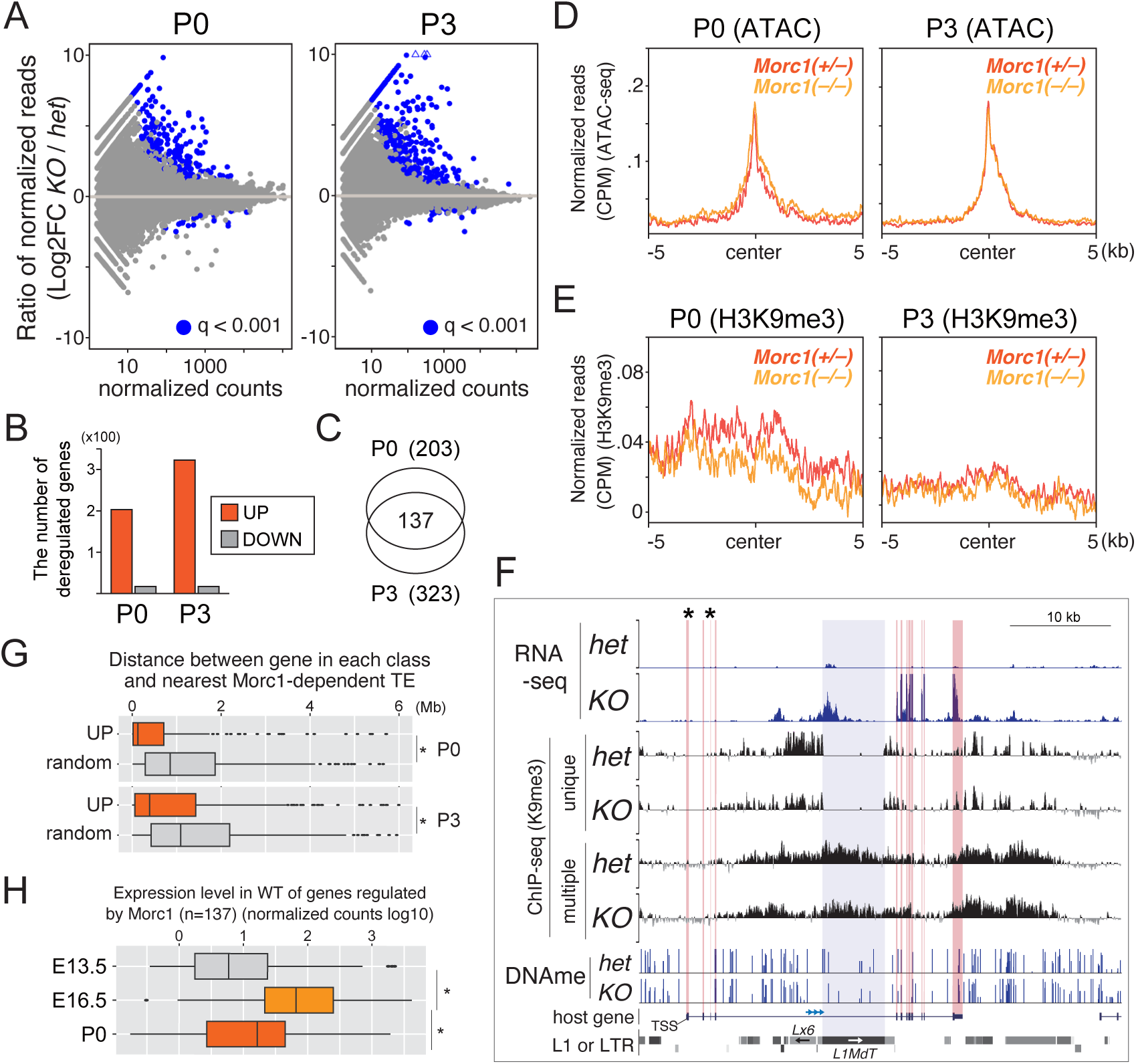
Morc1 reshapes the host gene transcriptome through suppression of neighboring TEs. (A) MAplots of RNA-seq reads mapped to host genes in *Morc1 KO* and *Morc1 hetero* gonocytes at P0 and P3. Blue dots denote differentially expressed genes (DEG) (p< 0.001). (B) Bar plots of the number of DEGs at the indicated timepoints. (C) Venn diagram showing overlaps of upregulated DEGs in *Morc1 KO* gonocytes between P0 and P3. (D) Line plots showing the average level of chromatin accessibility around TSS regions of upregulated DEGs in *Morc1 KO* gonocytes at the indicated time points in *Morc1 KO* and *Morc1 hetero* gonocytes. (E) As described in (D), but the H3K9me3 level is plotted. (F) Genome browser views of exons downstream from Morc1-dependent TEs inserted in the corresponding intron producing aberrant transcripts in *Morc1 KO* gonocytes. Asterisks indicate exons upstream from the same TEs. Note that such exons did not produce more transcripts in *Morc1 KO* gonocytes than in *Morc1 hetero* gonocytes. (G) Box plots showing the distance on chromosome between MdTE (ATAC) and upregulated genes (UP), downregulated genes (DOWN), and randomly selected genes (random). A t-test was applied for statistical hypothesis testing. *p < 0.01, **p < 0.05. (H) Box plot showing the abundance of transcripts of upregulated DEGs in *Morc1 KO* gonocytes using wildtype RNA-seq datasets at each developmental stage. A t-test was applied for statistical hypothesis testing. *p < 0.001.

**Figure 7.**
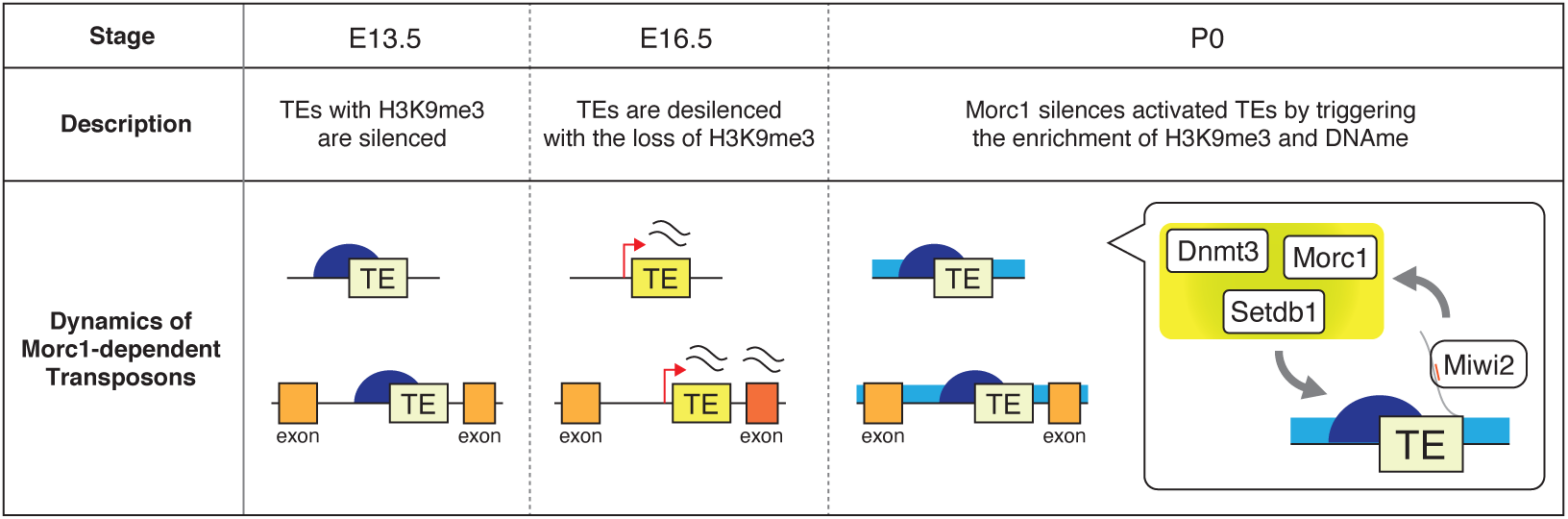
Model showing how Morc1 establishes the proper host transcriptome by suppressing transiently activated TEs during the gonocyte stage. From E13.5 to E16.5, TEs lose H3K9me3 from their TSS regions, leading to their transcriptional activation together with accumulation of their transcripts. As gonocytes enter the P0 stage, such TEs gain H3K9me3 and/or DNAme, and return to the transcriptionally repressed state. Morc1, SetDB1, and Dnmt3 family proteins are involved in establishing such an inactive chromatin state with the guidance of Miwi2.

## Discussion

### Morc1-dependent *de novo* formation of heterochromatin on TEs

In contrast to most reports, which reveal that epigenetic pathways maintain TE repression,^8^ this study focused on the mechanism of how to “establish” TE silencing during the mouse life cycle. In gonocytes, >10,000 copies of TEs are upregulated to possibly reorganize the overall chromatin structure, so that *de novo* DNA methyltransferases can gain access to the genome and catalyze DNAme along whole chromosomes.^35^ While this ectopic activation of numerous TEs would trigger chromatin reorganization, deregulation of TEs in general cause fatal genomic instability. Therefore, certain machineries re-silence active TEs. Here, we revealed that Morc1, which was previously shown to participate in DNAme on TEs, triggered enrichment of H3K9me3 and chromatin compaction over TEs in gonocytes. Target recognition of Morc1 was most likely mediated by the PIWI-piRNA pathway. Considering that only TEs actively producing transcripts were specifically silenced, the fact that small RNA-based targeting of active TEs operates in gonocytes is reasonable.

### Morc1 triggers both DNA methylation and H3K9me3 on TEs

Similar to mouse Morc1, AtMorc4 and AtMorc7 in *Arabidopsis thaliana* are involved in DNAme enrichment and TE silencing.^61^ AtMorc1 and AtMorc6 physically interact with HMTs for H3K9, SUVH2, and SUVH9.^43^ Moreover, the only MORC family protein in *C. elegans*, Morc-1, is involved in H3K9me3 enrichment in specific genomic regions.^62^ Additionally, mouse Morc2a and human Morc2 both have functional and physical interactions with the HUSH complex, which represses TEs through H3K9me3 enrichment.^44,45^ These data suggest that MORC family proteins use either DNAme or H3K9me3 to exert their repressive effect, depending on the biological context, although some evidence implies their ability to directly compact chromatin without any change in such epigenome marks.^58,63^ Compared with previous reports, it is noteworthy that mouse Morc1 triggers accumulation of both DNAme and H3K9me3 on TEs. While we showed that SetDB1 was a corresponding enzyme for H3K9me3 enrichment in gonocytes, the physical molecular network surrounding Morc1 has not been identified, which is an issue that should be addressed in the future.

### Small RNA pathway in Morc1-dependent chromatin modification

A previous report suggested that Morc1 and Miwi2 act separately in silencing TEs.^41^ This result was based on RNA-seq data from whole testes at P10, suggesting that the levels of transcripts from isolated germ cells at embryonic or newborn stages are different from the mixture of somatic and germ cells in testes. In our study using H3K9me3 ChIP-seq datasets, we observed a large amount of overlap between Miwi2 and Morc1 targets. In worms, Morc-1 acts in the RNAi pathway and plays a crucial role in germline immortality.^64^ Moreover, the H3K9me3 level on targets of HRDE-1, an Argonaute protein in worms, is reduced in Morc-1-mutant worms.^62^ These data support our model that mouse Morc1 exerts its repressive effect on TEs via a specific small RNA pathway.

### Complex interplay between DNA methylation and H3K9me3 on TEs in gonocytes

With the aid of Trim28, specific KRAB-ZFPs recruit SetDB1 to genomic regions of TEs, triggering their repression. Using a specific reporter containing the TE sequence, deletion of Trim28 or SetDB1 abolishes both H3K9me3 and DNAme.^56^ Similarly, G9a is required for DNAme on TEs in mESCs, indicating that H3K9me3 precedes DNAme in such processes.^57^ Conversely, we observed some clear differences between MdTE (K9me3) and MdTE (DNAme) at the family/subfamily level. This is consistent with results from mESCs, in which the overall level of DNAme at TEs was unchanged or only modestly reduced in *SetDB1 KO* cells.^65^ Moreover, DNAme around TSSs of host genes is only weakly correlated to H3K9me3 in somatic cells and mESCs.^66,67^ Therefore, it is conceivable that Morc1 induces accumulation of DNAme and H3K9me3 independently.

In summary, we found that TEs in gonocytes, whose expression was tolerated by the host system, were re-silenced by Morc1 at the chromatin level with accumulation of H3K9me3. Because *Morc1 KO* male mice show severe hypogonadism, this re-establishment of heterochromatin over activated TEs would ensure proper male fertility. Interestingly, host genes near Morc1-dependent TEs were activated at E16.5 in wildtype gonocytes (Figure 6H). Therefore, during this time window, new genes that are not transcribed from a normal TSS are expressed. The appearance of genes with a new domain conformation allows such genes to be domesticated by host system.^3,68^ In this regard, the gonocyte period could be a specific time window in which new genes emerge and evolve. Regardless, Morc1 plays a crucial role in coordinating interactions between host genes and TEs via epigenetic modification.

## Methods

### Animal care and use

All animal procedures were approved by the Institutional Safety Committee on Recombinant DNA Experiments and the Animal Research Committee of The University of Tokyo. Animal experiments were performed in accordance with the guidelines for animal experiments at The University of Tokyo.

### Generation of Morc1 knockout mice

*Morc1 KO* mouse was generated by introduction of Cas9 protein (317–08441; NIPPON GENE), tracrRNA (GE-002; FASMAC) and synthetic crRNA (FASMAC), and ssODN into C57BL/6J (CLEA Japan) fertilized eggs by electroporation. The synthetic crRNAs were designed to GCACTGGTTAAAAGGCCGTG (TGG) of the 1st intron of Morc1 and ATAAGGGACCAGATGAACAG (TGG) in the 20th intron. ssODN: 5’-GTGTTATTACTGGACACCAAGCAGATTCCACTGtttatttattGTGTGGAGGGCGGGTCACAAGAGAGATCGTGTG −3’ was used as a homologous recombination template. The electroporation solution contained 10 μM of tracrRNA, 10 μM of synthetic crRNA, 0.1 μg/μl of Cas9 protein and 1 μg/μl of ssODN in Opti-MEM I Reduced Serum Medium (31985062; Thermo Fisher Scientific). Electroporation was carried out by following previous reports.^69,70^ After electroporation, the embryos were cultured in KSOM^71^ medium for O/N, and then transferred into the oviducts of ICR (CLEA Japan) foster mothers at two-cell stage.

### Isolation of germ cells from testes

Testes were obtained from E13.5 and E16.5 Mvh-Venus TG embryos and newborn male Mvh-Venus TG pups.^72^ After removing the tunica, dissociation buffer [500 µl Dulbecco’s modified eagle medium (DMEM), 10 µl fetal bovine serum (FBS), 7.5 µl of 100 mg/ml hyaluronidase (Tokyo Kasei, Japan, H0164), 2.5 µl of 10 mg/ml DNAse (Sigma, D5025-150kU), 10 µl of 100 mg/ml collagenase (Worthington, CLS1), and 25 µl of 14000 U/ml recombinant collagenase (Wako, 036-23141)] was applied at 37 °C for 20 min. Subsequently, rigorous pipetting was performed until testicular cells were completely dissociated. After resuspending the cells in 2% FBS/PBS, Venus-positive cells were isolated by fluorescence-activated cell sorting using a FACS Aria III (BD). Before sorting cells, propidium iodine was added to select viable cells.

### ATAC-seq library construction

Using the original protocol,^73^ an ATAC-seq library was constructed from gonocytes as described previously.^35^

### ChIP-seq library construction

Venus-positive testicular germ cells (2×10^4^) were fixed with 1% formaldehyde for 10 min. Cells were resuspended in Swelling buffer [20 mM Hepes (pH 7.9), 1.5 mM MgCl_2_, 10 mM KCl, 0.1% NP-40, and 1 mM DTT]. After incubation on ice for 20 min, pelleted nuclei were resuspended in 1× shearing buffer (Covaris, 520154) and fragmented to ∼500 bp with a sonicator (BRANSON, SFX150). Fragmented products diluted with RIPA buffer [50 mM Tris-HCl (pH 8.0), 150 mM NaCl, 2 mM EDTA (pH 8.0), 1% NP-40, 0.5% sodium deoxycholate, and 0.1% SDS] were mixed by rotation with Dynabeads-ProteinG (Thermo Fisher, 10009D) or Dynabeads M-280 Sheep anti-mouse IgG (Thermo Fisher, 11201D) for 1 h at 4 °C. Immunoprecipitation was performed with 2 µl anti-H3K9me3 antibody (Active Motif, 39161) on Dynabeads-ProteinG (Thermo Fisher, 10009D) or 1 µl anti-H3 antibody (MBL, 16004) on Dynabeads M-280 Sheep anti-mouse IgG (Thermo Fisher, 11201D) overnight at 4 °C. Beads were washed with low buffer [0.1% SDS, 1% Triton X-100, 2 mM EDTA (pH 8.0), 150 mM NaCl, and 20 mM Tris-HCl (pH 8.0)] three times and then with high buffer [0.1% SDS, 1% Triton X-100, 2 mM EDTA (pH 8.0), 500 mM NaCl, snf 20 mM Tris-HCl (pH 8.0)] once. IPed products were eluted from beads by suspension in direct elution buffer [10 mM Tris-HCl (pH 8.0), 5 mM EDTA (pH 8.0), 300 mM NaCl, and 0.5% SDS] while shaking at 65 °C for 15 min. The sample was treated with proteinase K for 6 h at 37 °C, followed by reversal of crosslinking overnight at 65 °C. DNA was extracted by EtOH precipitation. Library preparation was carried out with a QIAseq Ultralow Input Library Kit (Qiagen, 180492) following the manufacturer’s instructions.

### ATAC-seq analysis

Reads trimmed to 25 bp were aligned to the mouse (mm10) genome using Bowtie2 with-N 1 and -X 2000 parameters, and uniquely aligned reads were extracted. After removing reads aligned to regions in the blacklist^74^ and PCR duplicates with Picard, we calculated Pearson correlation coefficients between 10 kb bins of biological replicates on chr1 using Deeptools. ATAC-seq reads were normalized to CPM. To calculate read counts on TE bodies, we also counted multiple mapped reads. Peak calling was performed using Homer with the following parameters: -style dnase, -size 500, and -minDist 1000. To define significantly accessible regions between *Morc1 hetero* and *Morc1 KO* gonocytes, peak call outputs and treated bam files were inputted to MAnorm. Significantly accessible regions were determined by the following criteria: ≥ 1 log2 fold change in normalized read counts and p-value < 0.0001.

### ChIP-seq analysis

Reads trimmed to 50 bp were aligned to the mouse (mm10) genome using Bowtie2 with the -N 0 option, and only unique reads were used for downstream analysis. After removing reads aligned to regions in the blacklist^74^ and PCR duplicates with Picard, we calculated Pearson correlation coefficients between 10 kb bins of biological replicates on Chr1 using Deeptools. ChIP-seq reads were normalized to CPM, and relative H3K9me3 enrichment was calculated by dividing ChIP enrichment of H3K9me3 by that of H3. To calculate read numbers on TE bodies, multiple mapped reads were included. Peak calling was performed using Homer with the following parameters: -style histone, -size 1000, and -minDist 2500. To define peaks that indicated significantly more or less H3K9me3 in mutant mice compared with wildtype mice, MAnorm and MAnorm2 were used. Significantly increased or decreased H3K9me3 region candidates were determined by the following criteria: ≥ 1 log2 fold change in normalized read counts and p-value < 0.0001 (calculated with MAnorm) or ≥ 0.8 log2 fold change in normalized read counts and p-value < 0.005 (calculated with MAnorm2). Among candidates, regions where fold changes of H3K9me3 read counts/that of H3 read counts were more than two were defined as significantly increased or decreased H3K9me3 regions. To construct line plots of the H3K9me3 level over the family/subfamily of TEs, we used mapping files that included multiple mapped reads.

### Genome enrichment calculation

RepeatMasker (open-0.4.5, mm10) was used to annotate TEs in the mouse genome (mm10). The proportion of each TE family was calculated for specific regions and the entire genome. The proportion in specific regions divided by that in the entire genome was defined as the enrichment value of Morc1 and Miwi2 targets. TEs occupying >0.1% of the TE population were included in the analysis. Values in figures were converted to logarithmic values.

### RNA-seq analysis

Reads trimmed to 90 bp were aligned to the mouse (mm10) genome using Hisat2 with the --dta option. After removing reads aligned to regions in the blacklist,^74^ we used FeatureCount to calculate read counts on TEs and genes with the following parameters: -p, -M (when calculating read counts on genes, -M was removed). Read counts were normalized to RLE using DESeq2. For gene annotation, we used mus_musculus_GRCm38_102.gtf (http://asia.ensembl.org/Mus_musculus/Info/Index).

### DNA methylome analysis

Datasets analyzed were deposited in Gene Expression Omnibus (GEO) under accession numbers GSE12757 and GSE63048, and in DDBJ under accession numbers DRP000638 and DRP002386. Raw reads trimmed to 50 bp were aligned to the mouse (mm10) genome using BSseeker2^75^ with default parameters. CGmaptools was used to calculate and visualize CG methylation levels on TEs.

### Organ culture

Organ culture was performed as described previously with some minor modifications.^55^ E16.5 testes without the epididymis were extracted from Mvh-Venus TG embryos and cut in half. DMEM (Nacalai) supplemented with 10% FBS (Gibco) and penicillin-streptomycin (Gibco) was used for the culture medium. Each testicular explant was placed in a 50 µl drop of the medium hanging on the lid of a culture dish and incubated at 37 °C for 3 days. The medium was changed every day. The SetDB1 Inhibitor was SETDB1-TTD-inhibitor (Target Mol, T9742) dissolved in DMSO and used at 10 µM as the final concentration.

### ChIP-qPCR

The starting material used for ChIP-qPCR analysis was 8×10^3^ cells. ChIP was performed as described above. Real-time quantitative PCR was performed using PowerUp SYBR Green Master Mix (Thermo Fisher) and the StepOnePlus real-time PCR system (Applied Biosystems). Primer sequences used for qPCR are listed in Supplementary Table 1.

### DNA gel electrophoretic mobility shift assay

The electrophoretic mobility shift assay was performed following a published method^76^ with some minor modifications. A pCAGGS plasmid expressing Flag-tagged GFP or Flag-tagged Morc1 was transfected into HEK293T cells. The cells were resuspended in buffer C (20 mM Hepes-KOH, 25% glycerol, 0.42 M NaCl, 1.5 mM MgCl_2_, 0.2 mM EDTA, 0.5 mM DTT, and 0.1% NP-40) and subjected to ultrasonication (SFX250, Branson). Samples were centrifuged to remove cell debris. The supernatant was incubated with ANTI-FLAG M2 Affinity Gel (Merck) for 4 h at 4 °C. The resin was washed three times with buffer C and then incubated with 3× FLAG peptide solution. The eluted fraction was applied to an Amicon column (3 KDa cutoff, Merck) to remove the 3× FLAG peptide. Purified proteins were incubated with 234, 40, or 20 bp DNA substrates in 20 mM Tris-HCl (pH 7.5), 50 mM NaCl, 1 mM MgCl_2_, 1 mM DTT, and 0.1 mg/ml BSA at 30 °C for 30 min. The reactions were resolved by 1.5% (for 234 bp DNA) or 2% (for 20 and 40 bp DNA oligos) agarose gel electrophoresis in TAE buffer (pH 7.5) at 4 °C for 2 h. DNA (234 bp) was detected by SYBR Gold staining (Thermo Fisher Scientific). X-Rhodamine-labeled DNA oligos were visualized using a Typhoon FLA 9500 scanner (GE Healthcare). The sequences of DNA substrates used in the electrophoretic mobility shift assay were as follows: 234 bp, 5ʹ-CCA AGT CGA CAA ACA GCT ATT GTT AAC CCC CCT CCA CCA GAG TAC ATA AAC ACT AAG AAG AGT GGG CGG TTG ACG AAT CAG CTG CAG TTC CTA CAG AGG GTT GTG CTG AAG GCC CTG TGG AAG CAC GGC TTC TCT TGG CCT TTC CAA CAG CCG GTG GAC GCC GTG AAA CTA AAG CTG CCT GAC TAT TAC ACC ATC ATA AAA ACC CCA ATG GAT TTA AAT ACA ATT AAG AAG CGG-3ʹ; 40 bp, 5ʹ-TTT TGT ATT ATC CTT ATA CTT ATT TAC TTT ATG TTC ATT T /36-TAMTSp/-3ʹ; 20 bp, 5ʹ-TAC ATT GCT AGG ACA TCT TT /36-TAMTSp/-3ʹ.

## Data availability

All data in this study have been deposited in the Gene Expression Omnibus (GEO) under accession number GSE235429.

## Acknowledgments

We thank all members of the Siomi laboratory for discussions and comments on this study. We also thank Tetsuji Kakutani (The University of Tokyo), and Raku Saito (The University of Tokyo) for critical reading of the manuscript. We thank Johji Nomoto and Shuji Ohshima for maintenance of the mouse strain. We thank Mitchell Arico from Edanz (https://jp.edanz.com/ac) for editing a draft of this manuscript. We also thank the One-Stop Sharing Facility Center for Future Drug Discoveries (The University of Tokyo) for FACS. This study was supported by Japan Agency for Medical Research and Development (AMED) Grant Numbers 21bm0704041h0003 (to S.Y.) and 22jm0210084h0003 (to S.Y.), MEXT KAKENHI Grant Number JP19H05466 (to M.C.S.), JSPS KAKENHI Grant Number 22K06338 (to H.N.), JSPS KAKENHI Grant Number 19K06616 (to S.Y.), the Takeda Science Foundation (to S.Y.), the NOVARTIS Foundation (Japan) for the Promotion of Science (to S.Y.), and the Astellas Foundation for Research on Metabolic Disorders (to S.Y.).

## Author contributions

Y.U., R.M., G.N., and S.Y. conducted biochemical analyses of *Morc1 KO* mice. Y.U., R.M., H. Narita, and S.Y. performed bioinformatic analysis. H. Nishihara. supervised the enrichment analysis of TEs. R.N. supervised the detection of ChIP-seq peaks. N.T. and K.A. established the *Morc1 KO* mouse. R.H. and Y.K. supervised *in vitro* culture of testes. S.Y. and M.C.S. conceived the project and designed the experiments. S.Y. and M.C.S. wrote the manuscript with input from all authors.

---

**Supplementary Figure 1.**
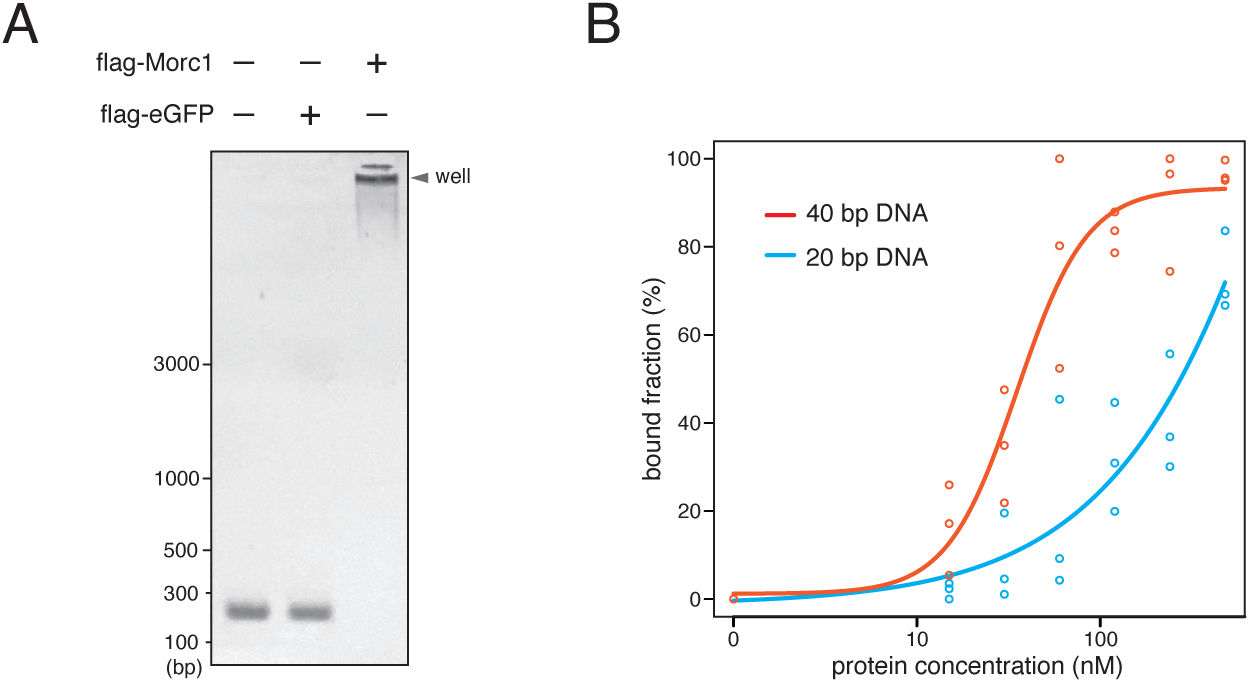
Morc1 directly binds to DNA without apparent sequence specificity. (A) Gel image of DNA electrophoretic mobility shift assays with or without the indicated proteins. The final concentration of each protein in the reaction buffer was 300 nM. Fourteen nanomoles of 234 bp linear dsDNA was used. (B) Quantification of gel shift analysis with 50 nM TAMRA-labeled 20 and 40 bp substrates. Protein concentrations tested were 15, 30, 60, 120, 240, and 480 nM. Three independent experiments are shown (circle). Lines are sigmoidal curves fit to the data.

**Supplementary Table 1 :**
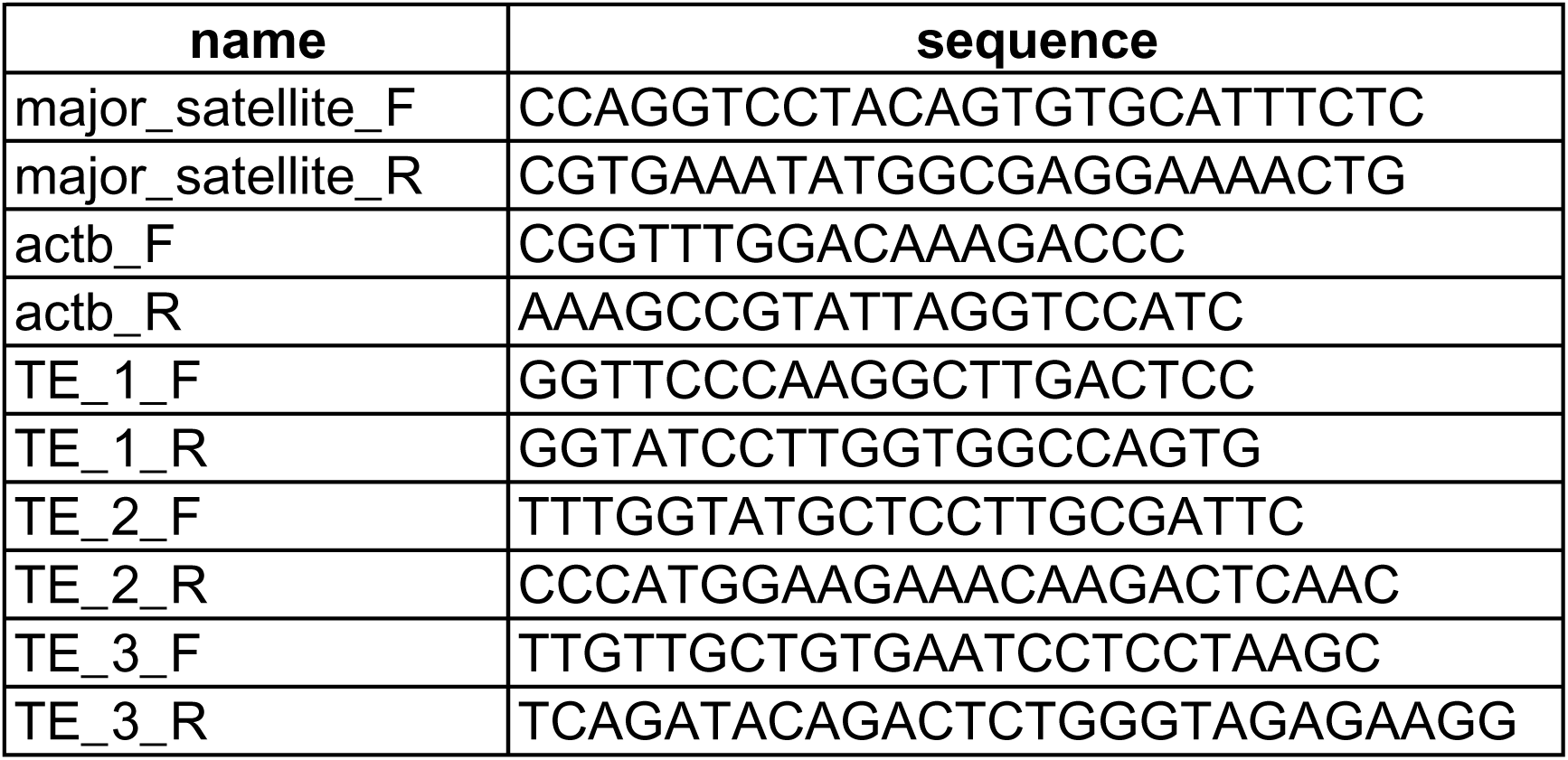
Sequence of primers used in this study.

